# The Genetic Puzzle of Multicopy Genes: Challenges and Troubleshooting

**DOI:** 10.1101/2024.03.25.586576

**Authors:** Vania Gabriela Sedano Partida, Henrique Moura Dias, Maria Teresa Portes, Marie-Anne Van Sluys

**Affiliations:** Departamento de Botânica, Instituto de Biociências, Universidade de São Paulo, 05508-090, SP, Brazil

## Abstract

**Background:** Studies with multicopy genes impose challenges related to gene redundancy and sequence similarity among copies. Recent advances in molecular biology and genomics tools associated with dedicated databases facilitate their study. Thus, the present work emphasizes the need for rigorous methodologies and standardized approaches to interpret RT-qPCR results accurately.

**Results:** The present work in *Physcomitrium patens* provides a comprehensive five-step protocol, using thiamine thiazole synthase (*THI1*) and sucrose 6-phosphate phosphohydrolase (*S6PP)* genes as proof of concept, to showcase a systematic workflow for studying multicopy genes. Beyond examining genes of interest, we highlight the critical role of choosing appropriate internal controls in the analytical process for accurately interpreting gene expression patterns. We emphasized the importance of identifying the most relevant orthologous gene, recognizing the inherent challenges in determining the most functional copy for subsequent studies. Our objective is to enhance comprehension of gene redundancy by dissecting multicopy genes’ genomic landscape and its characteristics. Furthermore, we address the decision-making process surrounding the quantification of expression levels of multicopy genes.

**Conclusions:** The study of multicopy genes discloses early events of functional adaptation. We emphasize the significance of multicopy genes in plant biology and provide a practical protocol for their study. Plant systems are strongly influenced by light/dark cycles, and the challenges inherent to this process are acknowledged. In conclusion, our comprehensive approach aims to advance the understanding of multicopy gene dynamics, offering practical methodologies and contributing valuable insights to the scientific community.

## BACKGROUND

Multicopy genes, or gene families, represent a fundamental aspect of plant genomes, contributing to the genomic complexity underlying plant species’ adaptability and resilience. These genes often arise through various duplication events, including whole-genome duplications and transposable elements activity, also playing pivotal roles in shaping the evolutionary trajectory of plant lineages (Flagel and Wendel, 2006; Fisher et al., 2014). In plant genomes, repetitive sequences, including multicopy genes, are particularly abundant, and constitute a significant portion of the genetic material (Kong, et., al 2023). Furthermore, multicopy genes frequently exhibit functional redundancy, which contributes to the robustness of plants facing environmental challenges. The evolutionary forces acting on duplicated genes may also lead to functional diversity, resulting in sub-functionalization or neofunctionalization, further enhancing the adaptability of plant species (Marakova, et., al. 2005). Multicopy genes play pivotal roles in plant biology, influencing plant development, stress responses, and adaptation to changing environments (Seidel et al., 2023). These genes are essential for shaping plant morphology and regulating growth (Stupar et al., 2006), also in response to external stimuli. Examples of multicopy gene families in plants include those involved in defense mechanisms against pathogens, response to abiotic stresses, and the biosynthesis of secondary metabolites (Weglöhner and Subramanian, 1994; Guitton and Berger, 2005). The existence of multiple copies within these families allows plants to fine-tune their responses to a dynamic and challenging environment.

The retention of duplicated genes is explained to promote neofunctionalization or sub-functionalization, which involves the acquisition of a new function or suggesting partitioning biochemical functions at the expression pattern level. Thus, without adverse environmental changes, the model theorizes similar expression levels among the paralogs. On the other hand, the absolute dosage and dosage-balance model also explains the retention of duplicated genes without a change in function (Makarova, et al. 2005; Qian and Zhang, 2014).

Despite their significance, studying multicopy genes remain challenging due to inherent complexities such as gene redundancy, where multiple copies perform similar functions, complicating the efforts to decipher the specific contribution of individual copies (for a review; Zeira and Shamir, 2019). Additionally, the high sequence similarity among copies can confound an accurate characterization and analysis (Teo et al., 2012). Distinguishing between individual gene copies and understanding their distinct functions becomes complex, hindering a comprehensive understanding of their roles in biological processes. Recent advances in molecular biology and genomics tools advancements have significantly enhanced our ability to study multicopy genes. Next-generation sequencing technologies allow high-throughput sequencing, enabling the characterization of entire gene families (Reis-Cunha et al., 2022; Namias et al., 2023). These technological tools provide researchers with the means to explore the complexity of multicopy gene families in details. In this context, many databases and tools dedicated to multicopy gene identification facilitate the discovery and retrieval of relevant information (Manni et al., 2021).

The study of multicopy genes requires rigorous protocols to ensure accuracy and reproducibility. Methodological choices, including experimental design and data analysis, significantly influence results interpretation, indicating the importance of standardized approaches. These protocols address challenges related to sequence similarity and gene redundancy. Established methodologies ensure reliable findings and advance our understanding of multicopy gene contributions to plant biology. Previous studies on multicopy genes in plants, such as Das and Bansal (2019), has provided valuable insights. However, significant gaps persist, particularly in deciphering the functional nuances of individual gene copies and their contributions to specific biological pathways and plant adaptation (Li et al., 2016; Dias et al., 2023). Further research in this area is essential for unlocking the full potential of multicopy genes in applications such as plant breeding, biotechnology, and crop improvement. Understanding the implications of multicopy gene variability could revolutionize the strategies for enhancing crop resilience, productivity, and adaptability to a changing environment. Addressing these knowledge gaps holds promise for the practical application of multicopy gene research in shaping the future of agriculture and plant biology.

Here, we highlight the importance of multicopy genes in plant genomes, seeking to facilitate their investigation by presenting a streamlined and curated protocol. Recognizing its inherent complexity, we provide a practical framework for researchers to navigate challenges such as gene redundancy and sequence similarity. Taking advantage of recent advances in molecular biology and genomics, coupled with dedicated databases and tools, our protocol aims to allow researchers to unravel the intricate expression pattern of individual paralogs amid multicopy genes. By emphasizing the importance of rigorous methodologies and standardized approaches through a case study of *S6PP* and *THI1* genes in the model plant *Physcomitrium patens*, our protocol would contribute to the reliability and reproducibility of findings in this field.

## CASE STUDY IN THE MOSS *PHYSCOMITRIUM PATENS*

Higher plants have a biological complexity that makes their study challenging at the genetic and metabolic level (Yuan et al., 2008). The need of understanding the biological and genetic functions of these organisms has led researchers to seek model plants for their study (Izawa and Shimamoto, 1996; Koornne and Meinke, 2010; Rensing et al., 2020). Thus, the study of bryophytes emerges to contribute to the understanding of how land plants evolved and adapted to life on soil. Mosses are structurally less complex than flowering plants, while sharing fundamental metabolic and physiological processes. Hence, relevant information on biological and genetic functions are also shared across many groups of Embryophytes (Naramoto et al., 2022).

*Physcomitrium patens*, formerly *Physcomitrella patens*, is a model organism due to its simple body-plant organization, large cells in both protonema and gametophore, as well as its small haploid genome 470 Mbp organized into 27 chromosomes (Lang, et al., 2018), and a life cycle of approximately 12 weeks (Schaefer and Zrÿd, 2001) under controlled conditions. The adoption of model organisms ensures that it has been extensively studied and that it only facilitates the investigation of biological phenomena, but that is not always the case (Rensing et al., 2020). *P. patens* imposes its own challenges, mostly due to genetic redundancy, as a result of ancestral events of whole-genome duplication (WGD) between 30 and 60 million years ago (Lang et al., 2017), which is an important eukaryotic evolutionary force resulting in the retention of a large number of duplicated genes, mainly involved in metabolism (Rensing et al., 2007).

*P. patens*, follows a life cycle that is typical of mosses and many other non-vascular plants. Its life cycle is referred to as “alternation of generations” and involves two distinct phases: the generation of multicellular haploid gametophytes alternating with a generation of morphologically distinct diploid sporophytes. The cycle begins with the germination of haploid (1n) spore, into a filamentous structure, characterizing the transient juvenile stage of the gametophyte, known as protonemata (Rensing et al., 2007). Protonemata consists of caulonema and chloronema cells. These filamentous structures serve as the juvenile stage of the plant and provide a surface for further development (Kofuji and Hasebe, 2014). The mature stage of the gametophyte, known as the gametophore, exhibits a heightened structural complexity featuring phyllodes (resembling leaves), stems, and rhizoids (root-like structures) (see Figure 1A). The progression from juvenile to the adult gametophyte is instigated by the differentiation of initial cells within the protonema filament, leading to the development of buds. The gametophore represents the adult haploid phase of the plant, serving for sexual reproduction. This process involves the initiation of a structure, the sporophyte, at the apex of the gametophore marking the transition to the diploid phase (2n), localized within the sporophyte. Upon maturation, the sporophyte releases spores (see Figure 1A), which, upon germination, give rise to new protonemata, thereby initiating a new cycle of the haploid phase in the plant’s life cycle.

Thus, the model plant *P. patens* was adopted as a case for investigation. In order to demonstrate the workflow potential proposed here for multicopy gene studies, we investigate the expression of the six thiamine thiazole synthase (*THI1)* (Dias et al., 2023a, 2023b) and five sucrose 6-phosphate phosphohydrolase (*S6PP)* (Partida et al., 2021) paralogs genes as a proof of concept. It highlights the significance and implications of identifying the appropriate orthologous genes among the multicopy gene family. This suggests that choosing the gene of interest is a critical decision. Challenges with multicopy genes may include determining which copy is functionally significant and understanding their roles. The experimental design focuses on two *P. patens* developmental stages: protonema and gametophore. In addition to two developmental stages, the experimental design considered the light/dark cycle over 48 hours. In total, 24 samples were collected in triplicates (Figure 1B). In the first stage, 1-week-old protonema cells are present, and in the second stage, 3-week-old adult gametophore phyllodes, stems and rhizoids predominate.

## STANDARDIZED APPROACH TO WORKFLOW DEFINITION

As mentioned above, a streamlined and curated protocol enables the understanding of intricate expression patterns of individual paralogs amid multicopy genes with reliability and reproducibility (Figure 2). Combining bioinformatics tools and experimental techniques ensures a thorough understanding of the genomic context and expression of multicopy gene families in diverse biological systems. Researchers can adapt and modify this protocol to suit specific organisms and research objectives, facilitating in-depth investigations into the role of multicopy genes in biological processes.

### STEP 1: Understanding the features of multicopy genes

Background: Identify multigene family members in a genome of interest.

Determining the significance of individual gene copies imposes a challenge, but our systematic investigation contributes substantially to unravel this complexity. Our objective is to enhance comprehension of gene redundancy by dissecting the genomic landscape and characteristics of multicopy genes and to proposing a protocol for RT-qPCR validation assays. Specifically, we selected multicopy gene families associated with sucrose 6-phosphate phosphohydrolase (S6PP), commonly known as SPP, and thiamine thiazole synthase (THI1) as target genes.

In the initial phase of the protocol, computational analyses play a central role in the meticulous identification of gene candidates. The Basic Local Alignment Search Tool (BLAST) (Altschul et al., 1990) was employed for a hypothesis-driven query, aligning it with distinctive features of multicopy genes under scrutiny. We chose the BLASTp algorithm from the Phytozome database, which maintains the *P. patens* v3.3 genome and relevant metadata annotation for putative subjects. Parameters were set rigorously to balance sensitivity and specificity based on E-value (> 1e-5), identity (> 70%), and coverage (> 70%).

An integral aspect of our strategy involves iterative refinement, wherein filtering strategies are applied to eliminate false positives and negatives, encompassing domain prediction (optimized by Yang et al., 2019). Domains representing conserved functional or structural units provide crucial insights into gene copy differentiation, evolutionary dynamics, and potential functional roles. The inclusion of domain annotation validates and improves the accuracy of homologs identification, introducing a new layer of information (see references Basu et al., 2019 and Wang et al., 2021). Through this process, we achieve complete identification and validation of S6PP (IPR006380) and S6PP-C domains (IPR013679) in S6PP orthologs, as well as the THI4 domain (IPR002922) in THI1 (refer to Figure 3A-B). We emphasize the importance of an integrative approach to multicopy gene identification. These parameters are generally sufficient for determining homologs and were applied to THI1 and S6PP. Following this parameter setup, we confirmed six THI1 homologs copies (Pp3c20_13540, Pp3c20_13770, Pp3c23_6510, Pp3c23_6600, Pp3c23_6580, Pp3c24_10800), located on chromosomes 20, 23, and 24, including tandem duplications of THI1 copies on chromosomes 20 and 23. Two excluded copies do not conform to features (Pp3c8_11240 and Pp3c22_8930) with coverages below 70% (see Figure 3A). While this approach has demonstrated to produce reliable results, it is important to highlight that its efficacy depends on the most conserved domains of the sequence (Albà and Caestresana, 2007; Kerfeld and Scott, 2011). In the search of S6PP homologs in *P. patens*, adjusted criteria are employed, considering an identity below 52% but with coverage exceeding 90% (refer to Figure 3B). The S6PP homologs Pp3c10_9450, Pp3c14_5810, Pp3c22_1840, Pp3c24_1340 contains S6PP-like and S6PP-C domains, supporting the presence on chromosomes 10, 14, 22, and 24. Pp3c19_6350 presents two incomplete domains, presumably this copy has lost its phosphohydrolytic function.

### STEP 2: Expression Profile by RNAseq Brings First Clues About Multicopy Genes

Background: make use of available RNAseq data to identify within the multicopy gene family which paralogs are expressed (also relevant to primer design)

After identifying potential homologs in Step 1, evaluating the coverage and expression profiles becomes essential to distinguish authentic and biologically relevant copies from pseudogenes. Authentic genes typically display distinct expression patterns that hold biological significance. Examining the expression profile allows the evaluation if the observed expression aligns with known biological functions. A consistent and contextually relevant expression profile supports the validity of a gene, while erratic or inconsistent patterns may indicate artifacts. To gain a preliminary view of the paralogous genes transcription, we explored the expression profile, which refers to the pattern of gene expression across different conditions or samples using the available PEATmoss database (Fernandez-Pozo et al., 2020). Examining how a gene behaves under various circumstances provides insights into its temporal and tissue-specific expression conditions (Figure 4).

Each of these datasets likely involves different experimental designs and conditions. This approach is beneficial because it allows the examination of expression patterns of the target genes in various contexts, potentially uncovering biological scenarios. Tissues or cell types may have specific gene expression patterns, and exploring these patterns can shed light on the expression of the gene paralogs in different parts of the organism. They often have similar functions but can present distinct expression patterns. Analyzing the expression of paralogs can provide insights into their functional divergence or redundancy. The initial analysis of gene expression using RNA-seq can serve as a foundation for designing future experiments. It provides a preliminary understanding of the gene’s behavior, and we can use this knowledge to guide more specific, hypothesis-driven studies.

While the expression profile provides valuable biological insights into a genés temporal and spatial profile, technically, distinguishing genuine expression from sequencing errors or artifacts becomes challenging when a gene exhibits low read coverage. Sufficient coverage is imperative to mitigate the probability of false positives or negatives (Sims et al., 2014). For instance, higher coverage enhances the confidence that the observed expression faithfully reflects the gene’s actual activity.

Our study revealed how valuable of using multiple RNAseq datasets to explore the expression patterns of paralogous genes (*S6PP* and *THI1*) under varying experimental conditions and tissue types. The results reveal that some paralogs are not expressed, revealing that not all copies are good candidates for studying function in the model organism. In addition, this approach can provide a comprehensive view of how these genes are regulated, supporting meaningful biological insights from the data. These findings highlight the dynamic nature of *THI1* and *S6PP* multicopy gene expression in response to different developmental stages and environmental conditions. The paralogs appear to have specific expression patterns based on the experimental context, with some being more prevalent in certain conditions or tissues. This information can be valuable for understanding the functional diversity of such gene copies and their relevance in different biological processes. Also, it is key to defining the primer design, as discussed below (STEP 4). It serves as a fundamental step for the RT-qPCR experiments, providing an overview of the selected tissues and stages for analysis.

### STEP 3: Crafting Experimental Design

Background: consider the organism’s biology and gene primary function to define the experimental design

Defining an experimental design and sampling time is crucial, as gene expression can vary over time (Liu et al., 2016) and across cell types. Different developmental stages, environmental conditions, and tissue types can significantly impact the expression of the target gene, yielding different results. The absence of expression doesn’t necessarily mean that the gene is non-functional. Genes can be silent under specific conditions or time points, even if they play crucial roles in other conditions. This implies that researchers must carefully consider these factors when designing their experiments and hypotheses (Huggett et al., 2005). Under this scenario and according to previous results (Partida et al. 2021; Dias et al. 2023), an experiment was set to investigate the transcriptional profile of *S6PP* and *THI1* paralogs in two synchronized stages of *P. patens* development (protonemata and gametophyte) and during the light/dark cycle. Plant material was sampled for 48 hours (every 4 hours) along the light/dark cycle on weeks 1 and 3 (Figure 1B). For this study, all copies of *THI1* and copies *Pp3c10_9450, Pp3c14_5810, Pp3c22_1840* and *Pp3c24_1340* of *S6PP* were selected based on the expression results of STEP 2 RNAseq analysis. Total RNA was extracted, and cDNA was prepared as described in the methods section.

Among the most common approaches to study gene expression patterns, it is the quantification of expression by the qPCR method (Bustin et al., 2009). This method often uses the selection of primers in conserved regions. However, in the case of multicopy genes with no previous knowledge of evolving functions, this strategy can lead to potentially incongruent conclusions. Thus, an essential step is the selection of specific primers that can distinguish the copies.

### STEP 4: Primer Design and References Definition

Background: uncover the reference (housekeeping) gene for the experimental design chosen RT-qPCR is a widely used technique for measuring mRNA expression levels due to its sensitivity and specificity. However, the success and accuracy of this method depend on various factors, including the sample quality, primer specificity, technique efficiency, and data analysis methods (Die, et al. 2010). Normalization is a fundamental step in RT-qPCR, and it is one of the most critical challenges. It involves selecting a housekeeping gene (also known as reference gene), that plays a critical role for normalizing gene expression data ensuring accurate and reliable results.

The choice of the reference gene significantly influences the accuracy of transcriptional quantification and further interpretation. To ensure reliable normalization, it is essential to choose a reference gene that exhibits stable expression across the conditions tested (Radonic et al. 2004; Huggett et al. 2005). For housekeeping genes, if instability is observed, may be necessary to reevaluate and select alternative internal controls guaranteeing that the selected reference genes remain stable across experimental conditions, crucial for accurate data normalization. If the selected reference gene demonstrates notable fluctuations in the expression level across the samples analyzed, it may result in misinterpretations capable of introducing bias and compromising the analysis accuracy. Researchers often underestimate the importance of rigorous reference gene selection. Pre-existing reference genes from the literature are sometimes used without validation in the specific experimental conditions. We emphasize that the choice of reference genes is a critical factor in gene expression analysis. It affects the reliability and validity of the results, and therefore, it is essential to assess the stability of potential reference genes in the specific experimental context to avoid incorrect interpretations of gene expression data. Therefore, it is crucial to assess the stability of potential reference genes specific to each experimental design, before applying RT-qPCR techniques for target gene analysis.

Given the significance of selecting a suitable reference gene, the expression of constitutive genes in P. patens was monitored adhered to the principle that the expression of transcripts should remain constant across the tested conditions and developmental stages, without exhibiting fluctuations in expression levels. We analyzed four commonly used housekeeping genes: PpE2, PpEf1α, PpST-P2α, and PpVH+PP (Le-Bail, Scholz and Kost, 2013), to define which would be the most adequate to our experimental design. The expression levels of each gene were analyzed in at least six time points that contemplate 24h cycle (light and dark) in the protonema development phase.

The statistical test of the 25th and 75th percentiles of the CT values for each gene, indicated that the PpE2 gene had the most stable expression among the four candidates evaluated (Figure 5). The choice of the appropriate reference gene depends on the specific experimental conditions to be tested. This study emphasizes the importance of rigorously evaluating and selecting an appropriate reference gene for RT-qPCR experiments to ensure the accuracy and reliability of gene expression data. It also highlights the context-dependent nature of reference gene selection, reinforcing the need to carefully consider the experimental design when making this choice.

The choice of the reference gene, being considered in the calculation of the quantification of gene expression, directly influences the biological interpretation of the results. If the reference gene shows fluctuations throughout the analyzed samples, the result can lead to an erroneous interpretation (Czechowski et al. 2005; Gutierrez et al. 2008; Bustin et al. 2009; Artico et al. 2010). Therefore, PpE2 is considered the most suitable housekeeping gene for this particular experimental design. The results support that even though there may be indications of suitable reference genes in the literature, there is no universal reference gene for a given organism.

### STEP 5: Selection of the modeling data

Background: Quantification of expression level of multicopy *THI1* and *S6PP* genes by RT-qPCR. Which to use: 2^-ΔCT^ or 2^-ΔΔCT^ equation, that is a relevant decision!

There are two types of expression quantification: absolute and relative. In the present work, we addressed the comparative method using relative quantification (Livak and Schmittgen, 2001), which assesses variations in the expression of genes (targets) within a specific sample (condition). This is achieved through normalization with a constitutive gene (housekeeping) or by examining changes relative to another reference sample (or condition), such as an internal control sample, guided by the 2^-ΔCT^ or 2^-ΔΔCT^ equations, respectively.

After normalization with housekeeping values, the 2^-ΔCT^ equation allows for comparisons among samples (or conditions) or between gene copies within a given sample (or condition). It is used for relative gene expression analysis within a single sample and does not provide information about the modulation of gene expression relative to other samples. It is a valuable tool for understanding gene expression levels in a specific context.

The 2^-ΔΔCT^ calculation is employed to assess alterations in gene expression concerning a control group. This method quantifies how many times a specific gene has been either upregulated or downregulated, being expressed as the fold change in comparison to a defined reference, referred to as an internal control sample.

Each equation will provide a specific information related to the expression of the target gene. To illustrate these approaches, let’s consider our case study involving two genes, THI1 and S6PP.The study utilizes 2^-ΔCT^ or 2^-ΔΔCT^ equations approaches to analyze their expression level under varying conditions. RT-qPCR assays were performed in both week 1 and week 3, along 48 hours and occurring under a light/dark cycle (16h/8h), as described in the experimental design section.

The 2^-ΔCT^ equation evidences that all copies of THI1 paralogs, both in protonemata tissues and adult gametophores, reached their maximum expression peak between 6PM and 10PM, which corresponds to the evening and early night. Conversely, the lowest expression is observed between 6AM and 10AM, corresponding to the early morning, suggesting a diurnal pattern of expression. It is noted that Pp3c23_6600 is more highly expressed in gametophore adult tissues, indicating that this specific paralog might play a more prominent role in the development or function of adult gametophores (figure 6A). The method, as described, doesn’t provide information about how the expression of the target gene is modulated, but it compares the expression of a gene across diferents conditions.

In the 2^-ΔΔCT^ approach, we studied the oscillation of gene expression at five time points; 2PM, 6PM, 10PM, 2AM and 6AM, during 48 consecutive hours, relative to 10AM that was used as an internal control in both protonema and gametophore for THI1. The analysis revealed that all THI1 paralogs exhibited rhythmic expression patterns over the 48-hour cycle in response to light/dark conditions, when compared to the 10AM control, with transcription levels increasing from 2PM and reaching their maximum peak of positive regulation between 2 PM and 10 PM. The modulation of expression levels differed between paralogs and developmental stages (protonema and gametophore). The maximum positive regulation for some *THI1* paralogs (Pp3c20_13540, Pp3c20_13770, Pp3c24_10800) occurred at 6 PM in protonemata, with significant fold changes, 75-fold, 200-fold and 20-fold, respectively (figure 6B). In gametophores, Pp3c23_6510 and Pp3c23_6600 had the highest expression levels at approximately 400-fold and 25-fold, respectively, but their modulation varied between protonemata. Expression modulation was not uniform across developmental stages, indicating distinct regulation profiles in protonemata and gametophores. Pp3c20_13540, Pp3c20_13770, and Pp3c24_10800 copies displayed rhythmic expression in both developmental stages. The five THI1 paralogs response to the light/dark cycle in protonemata tissues, while only three specific THI1 paralogs (Pp3c20_13540, Pp3c23_6600, and Pp3c24_10800) showed that regulation in adult gametophores (figure 6B). This suggests that their expression level are influenced by environmental cues, possibly related to light conditions and that the regulation of THI1 paralogs can vary between different tissues, with adult gametophores showing distinct patterns. The two equations support that THI1 paralogs are regulated by the light/dark cycle and that their expression patterns may be tissue-specific.

The use of the 2^-ΔCT^ equation to study the *S6PP* paralogs expression support its higher expression in protonema, with the maximum peak of expression between 10AM and 2 PM. We found that the Pp3c22_1840 copy is the most highly expressed among the S6PP homologs, as verified by RNAseq analysis, and it shows its maximum expression during the daytime period between 6AM and 6PM when there is light present in both protonema and gametophore. Conversely, the Pp3c10_9450 and Pp3c14_5810 copies show lower expression levels throughout the developmental cycle (figure 7A).

To analyse *S6PP* homologs, the 2^-ΔΔCT^ equation was applied with internal controls at 2 AM in protonema and 10 PM in gametophore, it was tested the same experimental setup as mentioned above for THI1 over a 48-hour light/dark cycle. The results showed, unlike *THI1*, there is no uniform modulation throughout the day among S6PP homologs. Pp3c10_9450, Pp3c14_5810, Pp3c22_1840 and Pp3c24_1340 homologs exhibited oscillation in modulation in both protonema and gametophore developmental stages (figure 7B).

All *S6PP* homologs are most expressed during the light period, from 6 AM to 6 PM. From 10 PM values decreased and reached their lowest fold change at 2 AM. Overall, the modulation of S6PP expression appears more pronounced in the adult gametophore phase when compared to protonema. Pp3c24_1340 homolog exhibited higher fold-change values than the control, indicating a greater change in the gametophore. Pp3c24_1340 is most expressed in the gametophore compared to protonemata. It reached a significant peak of approximately 70-fold at 6 PM in the gametophore. Pp3c10_9450 homolog had low fold-change values (ranging from 1 to 6-fold) in both gametophore and protonema. Pp3c14_5810 and Pp3c22_1840 copies showed the highest modulation peak at 2PM, with approximately 13-fold modulation and Pp3c10_9450 had a 7-fold increase at 10AM, all in protonema (figure 7B).

While the 2^-ΔΔCT^ quantification methodology revealed that the homologs are more modulated in the gametophore stage, we noticed that the expression levels are higher in young protonema tissues based on 2^-ΔCT^ quantification. These findings provide a detailed picture of how different *S6PP* homologs are expressed and modulated over a 48-hour light/dark cycle in different developmental stages. The data indicates that the expression patterns vary between homologs and between developmental stages, with some homologs exhibiting significant modulation during specific time periods of the day. The gametophore stage appears to have more pronounced modulation compared to protonema (figure 7B).

Overall, this information provides valuable insights into the regulation of THI1 and S6PP paralogs, their tissue-specific expression patterns, and their responsiveness to the light/dark cycle. These findings could have implications for understanding the biological processes and functions associated with these genes in the context of the organism’s development and environmental adaptation.

The two equations bring different biological information, the choice of calculation will always depend on the question that wants to be addressed and the way the information is approached. It’s important to note that 2^-ΔCT^ analysis is focused on the expression levels of the target gene in the given experimental design, and it doesn’t involve a comparison of modulation expression between different samples or conditions. This methodology assigns a value that provides biological information regarding the expression level of a gene in each condition and 2^-ΔΔCT^ calculation furnishes biological insights into the modulation of gene expression, utilizing a specific sample as a reference or control to discern the extent of modulation relative to it. In addition, the take home message of the present work is that the time of day when the sample is collected can interfere with the results obtained and eventually lead to a wrong biological interpretation.

## CONCLUSIONS

Our study addresses the challenges associated with multicopy gene analysis, emphasizing the importance of rigorous methodologies and standardized approaches. The provided protocol, exemplified in the model plant *P. patens* with *S6PP* and *THI1* genes, highlights the significance in the identification of appropriate orthologous genes. We focus on the variables that need to be considered when using gene expression quantification techniques, as well as differences in the evaluation of gene expression using different methodologies.

The streamlined and curated protocol presented here serves as a practical framework for researchers, utilizing bioinformatics tools and experimental techniques to understand intricate expression patterns of individual paralogs amid multicopy genes with reliability and reproducibility. Despite recent advances, significant gaps persist in deciphering the functional differences of individual gene copies and their contributions to plant adaptation. This approach can be adapted for specific organisms and research objectives, facilitating in-depth investigations into the role of multicopy genes in diverse biological processes. Further research in this area is crucial for unlocking the full potential of multicopy genes in applications such as plant breeding, biotechnology, and crop improvement, ultimately shaping the future of agriculture and plant biology.

## Acknowledgements

Authors acknowledge Dr Tatiana C. Pisetta technical support to maintain GaTELab infrastructure.

## Author contributions statement

VS and HMD conceived and outlined the experimental design, analyzed the data, prepared figures and tables, authored, reviewed and approved the final draft. MTP contributed with microscopy of *P. patens* life cycle, reviewed the manuscript, and approved the final draft. MAVS conceived the project, discussed results, authored, reviewed and approved the final draft.

## 2 SUPPLEMENTARY DATA LIST

Detailed material and methods description.

---

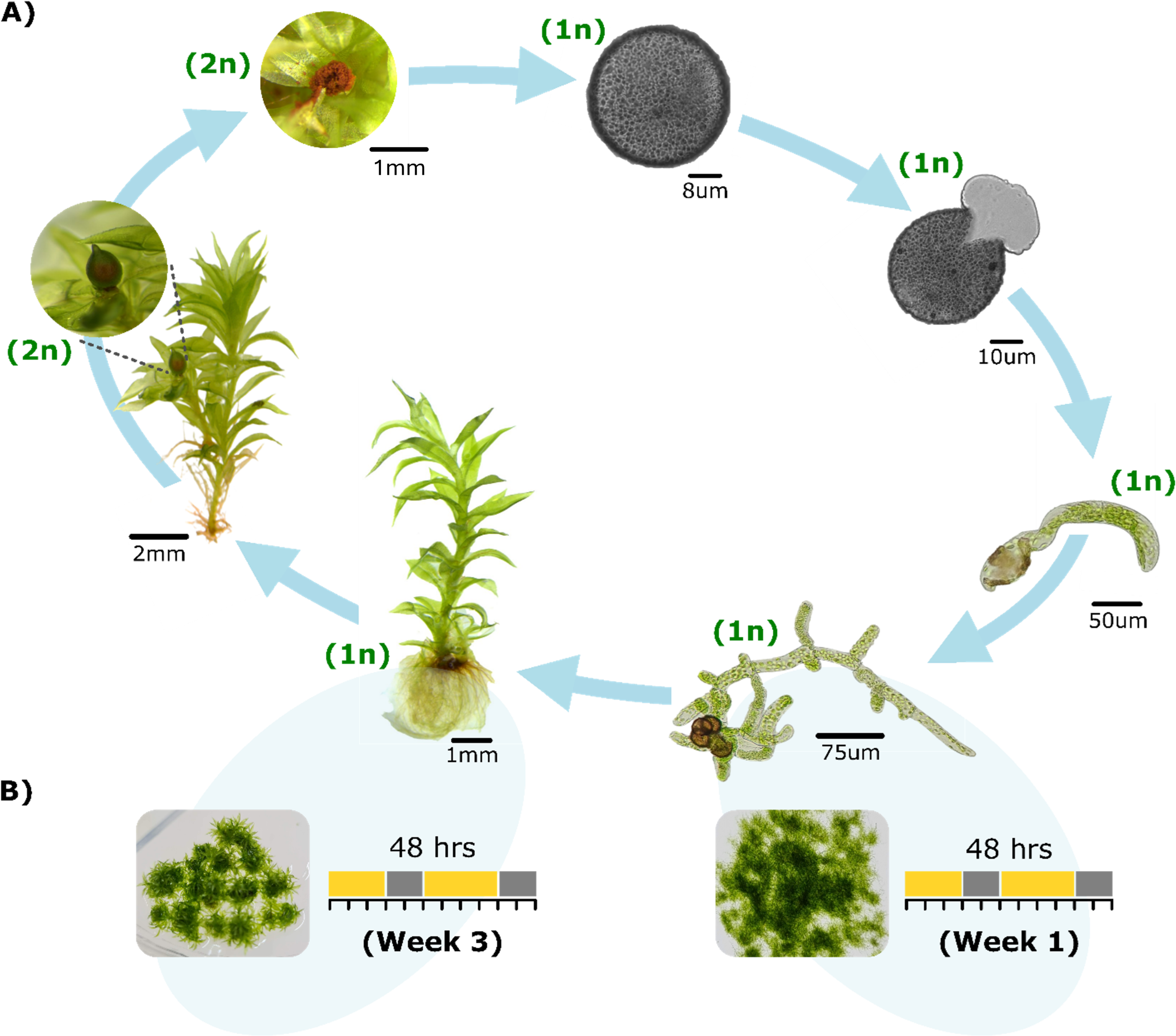

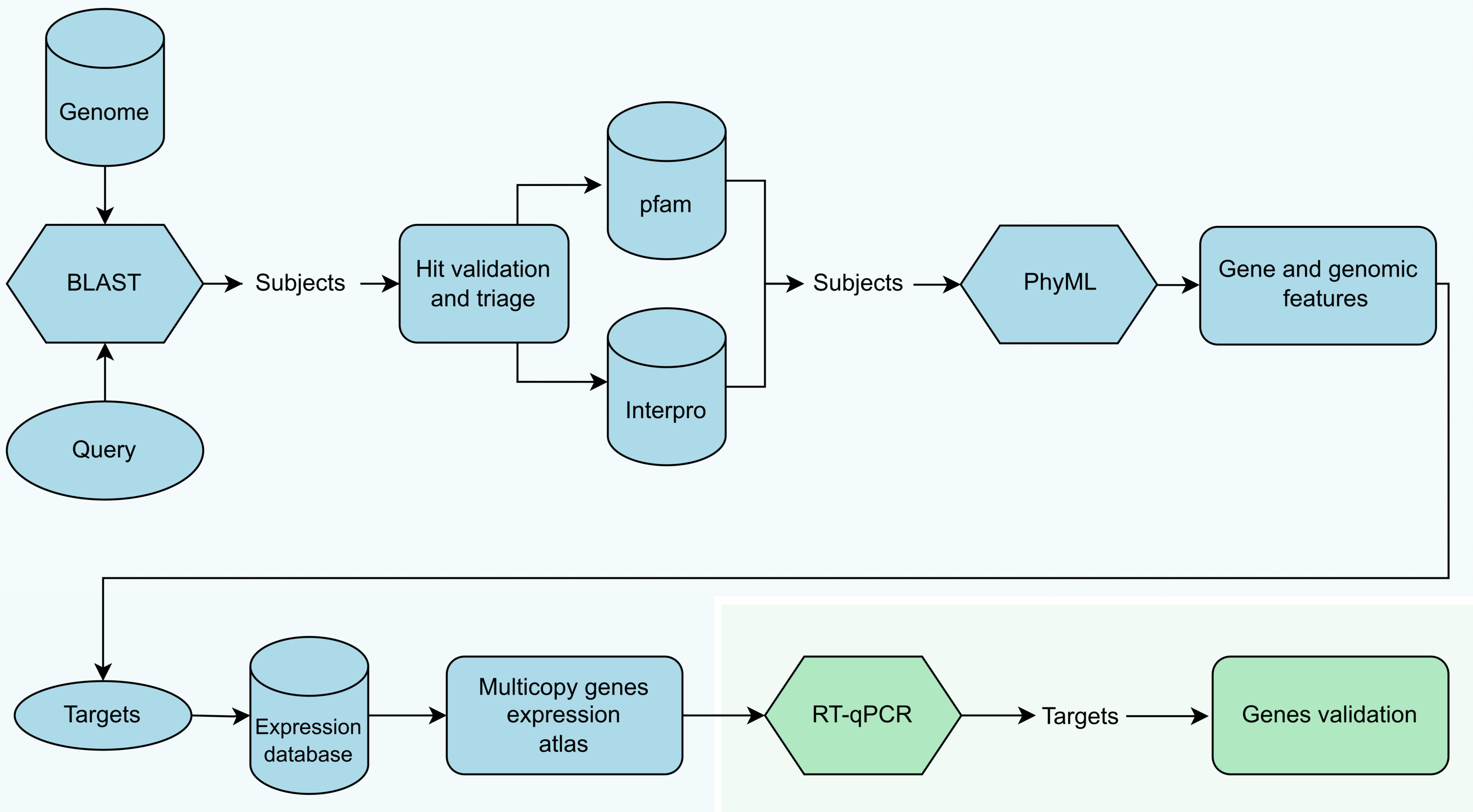

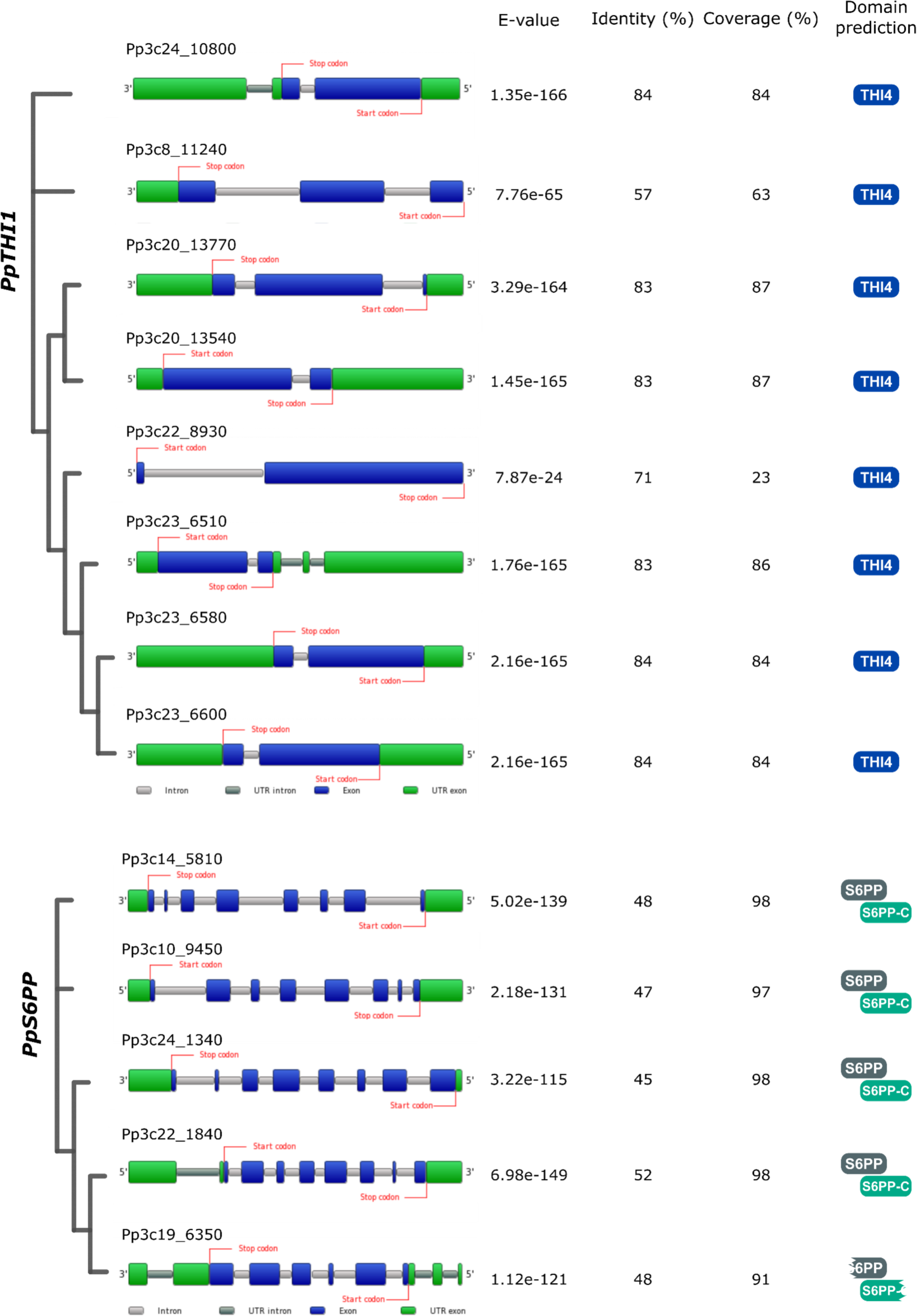

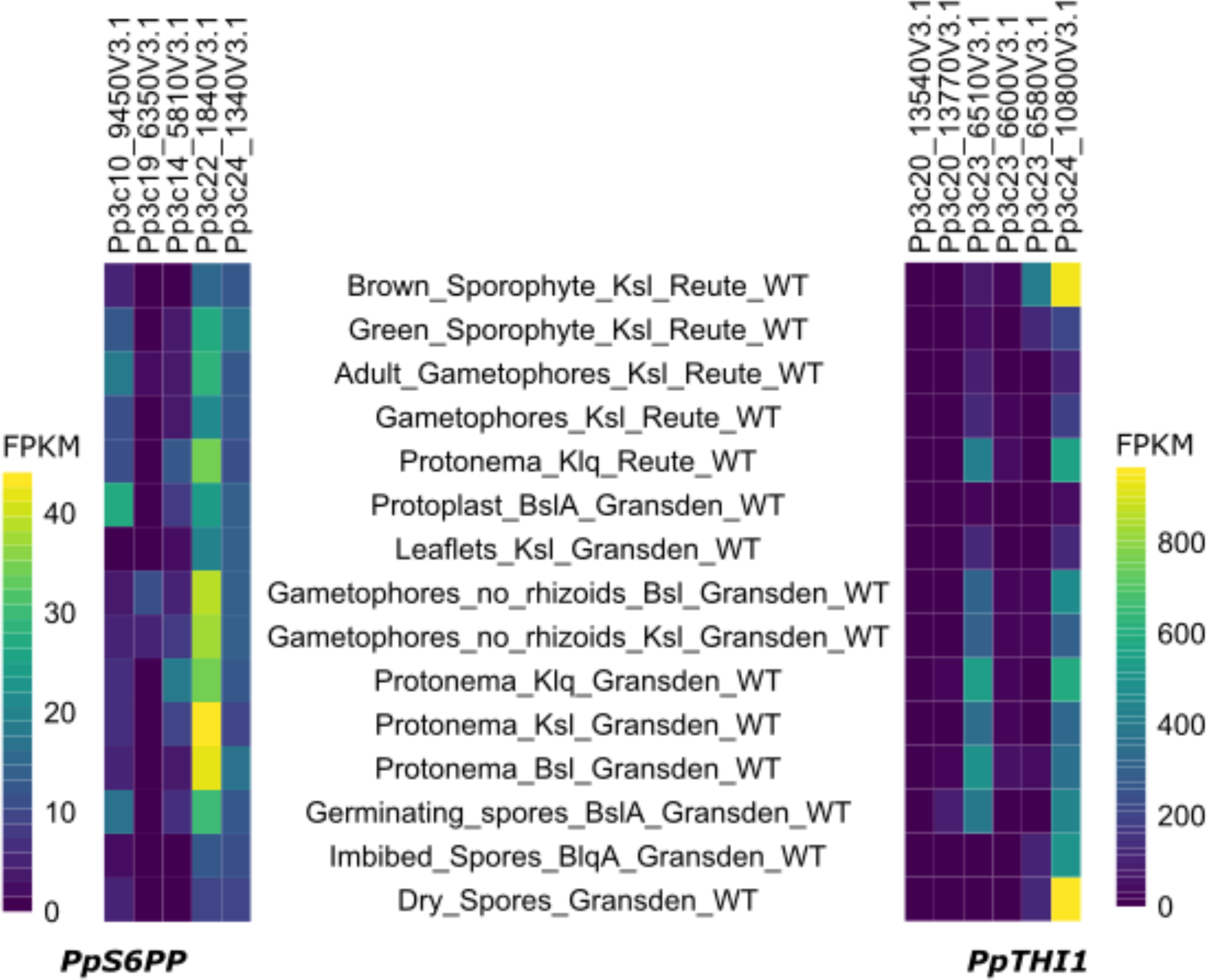

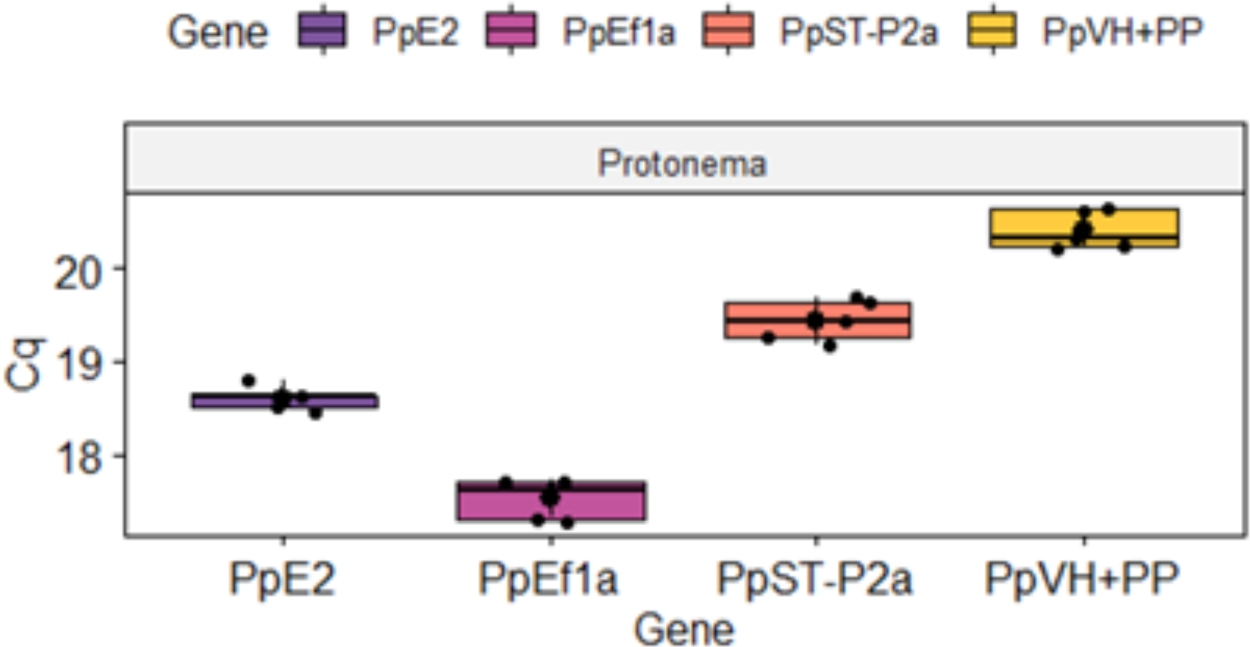

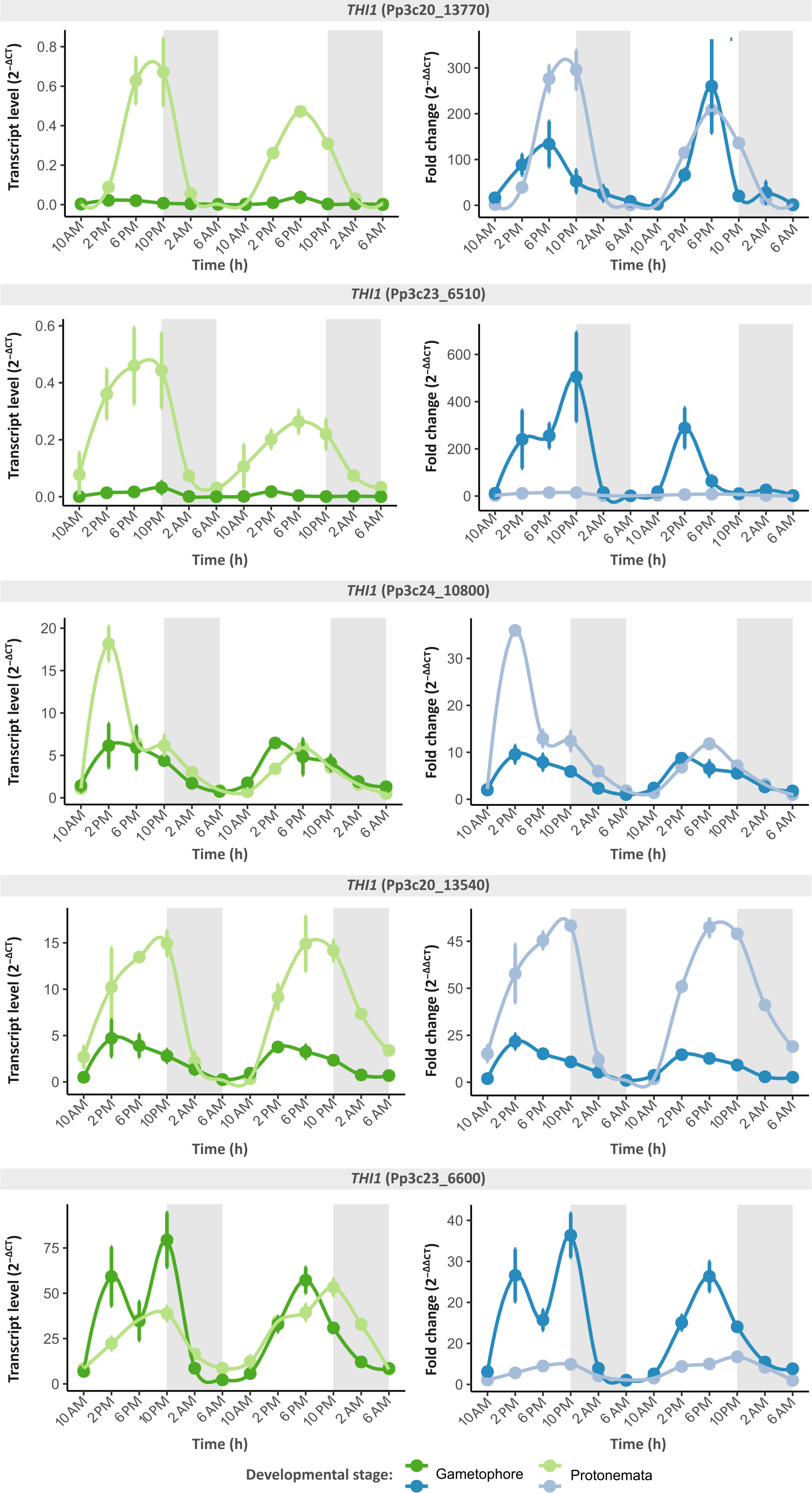

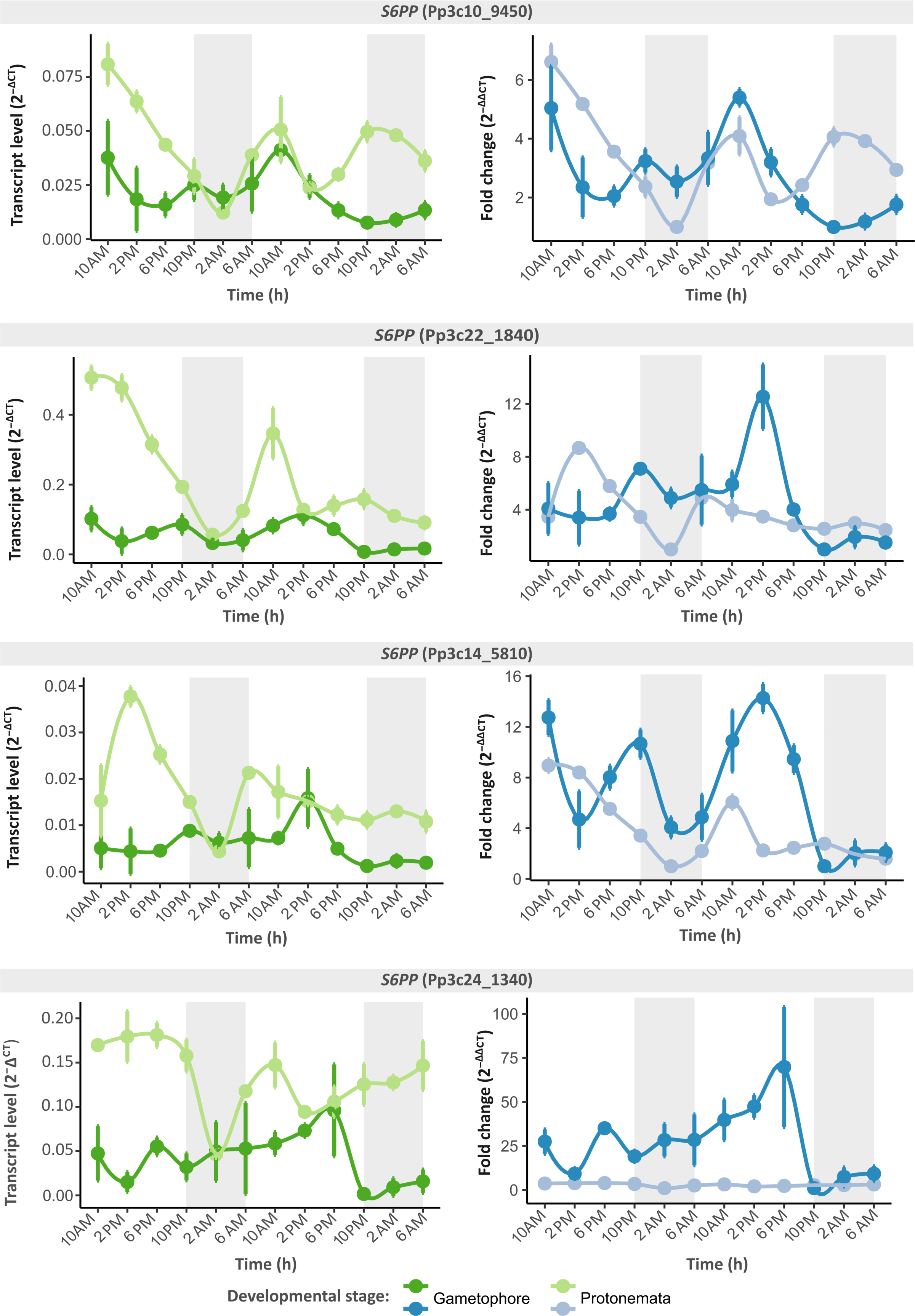

