## Supplemental Methods for "The Genetic Puzzle of Multicopy Genes: Challenges and Troubleshooting"

**Gene identification and genomic characterization**

The identification of multi-copy genes comprises a systematized method of sequential steps. Initially, (1) all protein datasets of the selected organism of interest undergo comparison through BLAST, utilizing an E-value cutoff of 1e-5, ensuring a minimum coverage of 70% for the query protein and an identity of at least 70% referencing a standard. Subsequently, (2) sequences meeting these criteria for both coding sequences (CDS) and proteins were systematically gathered. Following this step, the protein sequences are aligned using MAFFT v77.450, employing default parameters (Katoh and Standley in 2013). The alignment output serves as input for the subsequent step. To further validate the putative candidate genes, (3) a verification step is performed utilizing the online software InterProScan (Blum et al., 2021) and Pfam (El-Gebali et al., 2019) and (4) an additional sequence clustering is executed employing PhyML 3.0, with a conservative approach that considers bootstrap values (Guindon et al., 2010).

**RNAseq analysis**

To obtain an initial understanding of the transcription of paralogous genes, we examined the expression profile across various conditions or samples utilizing available RNAseq data. This step ensures that expressed paralogs within the multicopy gene family are identified, which is a crucial step for primer design.

Mapping was conducted using the Salmon software (Patro et al., 2017), accessible on the Galaxy platform (https://usegalaxy.eu/). Default mapping parameters were employed, and a BAM file was generated to visualize the coverage of the mapped reads. Visual verification of the mappings was performed using the Unipro UGENE software to confirm the comprehensive coverage of the transcripts (Figure S1).

**Plant Material and Growing Conditions**

Wild gametophyte ecotype Gransden 2004 of 30 days was fragmented with a sterile blender to return to the stage of cellular undifferentiation, the resulting blend was used as inoculum to start the growth of the material for the experiment. Initial inoculation was performed on plates with BCDAT medium. The growth and development of the inoculums took place in a phytotron with controlled conditions; photoperiod 16 hours of light (from 06:00 AM to 22:00 PM) and 8 hours of darkness (from 22:00 PM to 06:00 AM), temperature of 25°C, humidity 65% and light intensity of 150 µmol. The whole experiment lasted 3 weeks, where on week 1 protonema was the major tissue while on week 3 most of the cells already differentiated to gametophore and rhizoids (Figure 1). Sampling was carried out every 4 hours, being on schedule; 10:00 AM, 14:00 PM, 18:00 PM, 22:00 PM, 02:00 AM and 06:00 AM, for 48 consecutive hours, in the two mentioned stages. The experiment included three biological replicates for each time point. A total of 72 samples/conditions integrates the light/dark cycle analysis.

**RNA Extraction and cDNA Synthesis**

Each sample was ground to a fine powder and kept at -80°C until use. Total RNA was isolated employing TRIzol® reagent (Invitrogen®), following manufacturer instructions and digested with DNAse TURBO DNA-free (Ambion®, Life Technologies™) to remove any contaminating genomic DNA. Total RNA was visualized by agarose gel, to analyze integrity. 1000 ng of total RNA was used as cDNA template with reverse transcriptase SuperScript™ III (Invitrogen®), and oligo dT primer strategy, following the manufacturer's instructions and stored at -20°C until used.

**qPCR experimental design**

**Defining a housekeeping gene to the experimental design**

This study emphasizes the importance of rigorously evaluating and selecting an appropriate reference gene for RT-qPCR experiments to ensure the accuracy and reliability of gene expression data. It also highlights the context-dependent nature of reference gene selection, reinforcing the need to carefully consider the experimental design when making this choice. The choice of the reference gene, for the quantification of gene expression, directly influences the biological interpretation of the results. If a fluctuation of the reference gene is observed throughout the analyzed samples, the result can lead to an erroneous interpretation (Czechowski et al. 2005; Gutierrez et al. 2008; Bustin et al. 2009; Artico et al. 2010).

Oligonucleotides were designed for the amplification of constitutive genes, previously selected based on the literature (Le-Bail, Scholz and Kost, 2013), *ubiquitin-conjugation enzyme E2 (PpE2)*, *elongation factor 1α* (*PpEf1α),* *serine threonine protein phosphatase 2α* (*PpST-P2α)* and *v-type h(+) translocating pyrophosphatase* (*PpVH+PP)* were tested to choose the best housekeeping for the specific experimental design. The expression levels of each gene were analyzed in at least 6 sample points that represents the 24h cycle (light and dark) in the protonema developmental phase.

The RT-qPCR reaction was performed with the kit SYBR™ Green PCR Master Mix, by ThermoFisher Scientific, following the manufacturer's instructions with 0.3 µM of specific primer. CTs and dissociation curves were analyzed using the Design & Analysis 2.6.0 Real-Time PCR System, Applied Biosystems program from Thermo Fisher Scientific.

**Specific Primers Design for RT-qPCR Assay**

The primers were carefully designed to distinguish each copy, with at least one of the primers siting at an exon-exon junction. Primer sets, including housekeeping primers were verified in Primer Express™ v3.0.1 software. The oligonucleotides designed are shown in Table S1.

**Gene expression quantification by RT-qPCR assay**

RT-qPCR was assayed in both week 1 and week 3 along 48 hours each sampling, under light/dark cycle (16h/8h). The reaction was performed with the SYBR™ Green PCR Master Mix, by ThermoFisher Scientific with 0.3 µM of specific primer.

CTs and dissociation curves were analyzed using the Design & Analysis 2.6.0 Real-Time PCR System, Applied Biosystems program from Thermo Fisher Scientific. Gene expression was calculated using the 2^-∆CT^ or 2^-∆∆CT^ equation, considering CTs values of PpE2 gene selected as the housekeeping control.

**Analysis visualization methods**

Using the 2^-∆CT^ equation, gene expression levels are determined by normalizing the data using a reference gene, referred to as a "housekeeping gene". The expression level of the target gene is calculated relative to the housekeeping gene within a particular sample (condition). This means that the analysis produces a value that indicates the relative expression of the target gene normalized to the expression of the housekeeping gene in that sample (condition). With this approach, expression levels across all normalized samples can be compared within the context of the specified experimental design.

The calculation is guided by the equation:

∆CT = CT[Target]-CT[Housekeeping]

Expression level = 2^-∆CT^

The 2^-∆∆CT^ calculation is employed to assess changes in gene expression modulation. This involves normalization with a housekeeping gene and subsequent utilization of an internal control. By designating an experimental condition as a reference within the experiment, the expression levels are quantified as a fold change to the reference. This fold change indicates the relative increase or decrease in the target gene's expression in a specific sample (condition) compared to the reference sample (condition). The values can be interpreted in a biologically meaningful manner; a fold-change >1 indicates positive regulation, reflecting an increase in gene expression, and <1 is a negative regulation, signifying a decrease in gene expression.

The equation is:

∆CT = CT[Target]-CT[Housekeeping]

∆∆CT = (∆CT)- (∆CT [Control])

Fold-change = 2^-∆∆CT^ (Livak and Schmittgen, 2001)
